# A new-engineered integrative tool to target the terminal compartment of the *Streptomyces* chromosome

**DOI:** 10.1101/2025.02.27.640095

**Authors:** Nicolas Delhaye, François R. Pelé, Hoda Jaffal, Stéphanie Bury-Moné

**Author notes:** Address correspondence to Stéphanie G. Bury-Moné.

## Abstract

Phages are a valuable resource for the genetic engineering of *Streptomyces* antibiotic-producing bacteria. Indeed, a few integrative vectors based on phage integrase are available to insert transgene at specific genomic loci. Chromosome conformation captures previously demonstrated that the *Streptomyces* linear chromosome is organized in two spatial compartments: The central compartment encompassing most conserved and highly expressed genes in exponential phase, and the terminal compartments enriched in poorly conserved sequences including specialized metabolite biosynthetic gene clusters. This study introduces a new integrative tool based a recently described phage, Samy, which specifically targets the terminal compartment of its native host chromosome. Samy is related to PhiC31 phage and, like this latter, encodes a serine integrase. Whereas PhiC31 targets a site generally located near the origin of replication, the Samy integration site is one of the farthest known *attB* sites from it. We demonstrated that the Samy integrase efficiently mediates the specific integration of a non-replicating plasmid in six *Streptomyces* strains from distinct clades. Bioinformatic analyses revealed that the Samy *att*B site is rather conserved, and located in the terminal compartment of most *Streptomyces* chromosomes. Unusually for a serine integrase, Samy-based integration system exhibits slight excision. The Samy orthogonal site-specific recombination system extends the range of genetic tools available for engineering the *Streptomyces* chromosome by broadening the range of targeted locations, especially within the terminal spatial compartment.

**Key points:**

*-* Samy-based integrative vectors are new tools for engineering *Streptomyces* strains.
*-* They are among the few targeting the terminal compartment of the chromosome.
*-* Their integration is farthest from the replication origin in most strains compared to other vectors.

## Introduction

*Streptomyces* are renowned for their prolific production of specialized metabolites, including antibiotics, pesticides or pigments, with applications in medicine, agriculture and food industry. These bacteria are also well-known for their biodegradation efficiency, driven notably by the secretion of potent enzymes. Therefore, their genome engineering is therefore of great biotechnological interest (Lee et al. 2019; Mitousis et al. 2020). Temperate phages and integrative plasmids serve as crucial tools for the site-specific, stable, and possibly multiplexed insertions (Li et al. 2017; Li et al. 2019; Gao and Smith 2021) of (over)expression cassettes into genomes. This capability is essential for the long-term, large-scale biotechnological exploitation of *Streptomyces* strains. Orthogonal integration systems for *Streptomyces* genomes have been developed, based on tyrosine- recombinases [pSAM2 (Boccard et al. 1989; Raynal et al. 1998), VWB (Van Mellaert et al. 1998), µ1/6 (Farkašovská and Godány 2012)] or serine-recombinases [PhiC31 (Bierman et al. 1992; Groth et al. 2000), PhiBT1 (Gregory et al. 2003), TG1 (Morita et al. 2009), R4 (Miura et al. 2011), SV1 (Fayed et al. 2014), PhiJoe (Fogg et al. 2017), PhiOZJ (Ko et al. 2020), PhiWTR (Ko et al. 2020)]. These enzymes also named ‘integrases’ catalyze the specific recombination between *att*P (attachment site on the phage or plasmid) and *att*B (attachment site on the bacteria) sequences, generating *att*L (attachment site on the left) and *att*R (attachment site on the right) sites on either side of the integrated construct [for reviews, see (Merrick et al. 2018; Kormanec et al. 2019; Mitousis et al. 2020; Ba et al. 2023; Smith 2015)]. Site-specific recombination systems based on serine-integrases are considered as the simplest and more likely to function across a broader range of cells due to the absence of host factor requirement (Merrick et al. 2018; Ba et al. 2023). The integration is generally directional, meaning that the reverse reaction (*att*L x *att*R recombination) occurs only in the presence of specific partners. The PhiJoe and PhiBT1 integrases are exceptions to this rule since they demonstrate significant excision activity even when expressed alone (Zhang et al. 2008; Fogg et al. 2017).

The *Streptomyces* chromosome is linear - which is unusual for bacteria - and genetically compartmentalized into a central region harboring the origin of replication and cores genes, and two extremities populated by poorly conserved sequences (Bury-Moné et al. 2023). These extremities are enriched in genomic islands including specialized metabolite biosynthetic gene clusters (SMBGCs). The majority of these poorly conserved sequences remain transcriptionally silent and/or weakly expressed under lab conditions (Van Bergeijk et al. 2020; Lioy et al. 2021). As reported in *Streptomyces ambofaciens* ATCC 23877 (Lioy et al. 2021) and *Streptomyces coelicolor* A3(2) (Deng et al. 2023), the genetic compartmentalization correlates with chromosome architecture and gene expression: The distal ribosomal RNA (*rrn*) operons delimit a highly structured and expressed region termed ‘central compartment’, presenting structural features distinct from those of the terminal compartments which are almost transcriptionally quiescent during vegetative growth. This architecture and gene expression pattern are dynamic during over the *Streptomyces* developmental cycle (Lioy et al. 2021; Szafran et al. 2021; Deng et al. 2023).

The central compartment is considered as a hotspot for integrative element insertions (Choufa et al. 2022). In line with this observation, a systematic analysis of the location of remnant and complete prophage sequences along *Streptomyces* chromosome revealed that they are most commonly found in the half that includes the origin of replication (Sharma et al. 2023). Accordingly, the *Streptomyces* integrative vectors generated so far target the central compartment in most strains. These include site-specific recombination systems based on pSAM2 (Boccard et al. 1989; Raynal et al. 1998), PhiC31 (Bierman et al. 1992; Groth et al. 2000), VWB (Van Mellaert et al. 1998), PhiBT1 (Gregory et al. 2003), TG1 (Morita et al. 2009), µ1/2 (Farkašovská and Godány 2012), SV1 (Fayed et al. 2014), PhiJoe (Fogg et al. 2017), PhiOZJ (Ko et al. 2020) and PhiWTR (Ko et al. 2020), with the three last targeting the same *att*B site. The notable exception to this is the R4-integrase-based plasmids (Miura et al. 2011), which target a site located in the terminal spatial compartment of *Streptomyces* chromosome in most strains (Lorenzi et al. 2022).

We recently identified a new temperate phage named Samy, which is closely related to PhiC31 (Jaffal et al. 2023). Samy specifically targets the terminal compartment of the native host chromosome in *S. ambofaciens* ATCC 23877. While its prophage form is mainly silent under most growing conditions analyzed so far, the identification of its induction conditions revealed that the Samy productive cycle promotes the *in vitro* dispersal of *Streptomyces* multicellular structures in response to stress (Jaffal et al. 2023). In this study, we characterized a site-specific recombination system based on Samy serine-integrase. This expands the genetic toolkit for engineering the *Streptomyces* chromosome by broadening the range of targeted locations. To date, this new tool is capable of targeting the most distant site within the terminal compartment of most *Streptomyces* strains.

## Materials and Methods

### Strains, plasmids and growth conditions

The strains and plasmids used in this study are listed in **Table S1**, while all primers are provided in **Table S2**. *Streptomyces* strains were grown at 30°C on solid soy flour-mannitol (SFM) medium (20 g/L organic soy flour, 20 g/L mannitol, 20 g/L agar) (Kieser T. 2000) or in liquid Tryptone Soya Broth (TSB, 30 g/L Tryptic Soy Broth BD™ 211825) (Kieser T. 2000). The 2TY (16 g/L Bacto Tryptone, 10 g/L Yeast Extract, 5 g/L NaCl) and SNA (8 g/L Bacto Nutrient Broth) media (Kieser T. 2000) were used exclusively for conjugation. When appropriate, apramycin (50 μg/mL) and/or nalidixic acid (50 μg/mL) were added to the growth media. *Escherichia coli* strains were grown at 37°C in Luria-Bertani (LB) medium, supplemented with 50 μg/mL apramycin and/or 25 μg/mL kanamycin when appropriate.

### Vector cloning

Standard techniques were used for recombinant DNA manipulation. BsaI sites were introduced into pOJ260 by PCR amplification using SBM375 and SBM376 primers, generating pOJ260-MoClo. In parallel, a PCR product corresponding to the 56,510–58,494 genomic region of Samy phage (OR263580.1), including the *SAMYPH94* gene encoding Samy-integrase, its promoter, and the Samy-*att*P site sequences flanked by BsaI sites, was obtained by amplifying the Samy genome with SBM560 and SBM561 primers. The Samy genome was isolated from the supernatant of *S. ambofaciens* ATCC 23877 grown in BM (bacteriophage medium), as previously described (Jaffal et al. 2023). This insert was then cloned into pOJ260-MoClo via Golden Gate Assembly (NEBridge® Golden Gate Assembly Kit BsaI-HF® v2) according to the manufacturer’s recommendations.

A pOSV vector from the Aubry *et al*. collection (Aubry et al. 2019) conferring apramycin resistance was digested with AflII and SbfI. In parallel, a PCR product corresponding to the 56,510–58,494 genomic region of Samy phage (OR263580.1), flanked by AflII and SbfI sites, was obtained by amplifying the Samy genome with SBM588 and SBM589 primers. After purification and digestion with AflII and SbfI, both the vector and insert were heat-inactivated and ligated using T4 DNA ligase (PROMEGA).

### Conjugation

Plasmids intended for conjugation into *Streptomyces* were first introduced by electroporation into *E. coli* ET12567 carrying pUZ8002 to provide the necessary transfer functions. Conjugation was then performed as previously described (Kieser T. 2000). Briefly, 10⁸ *Streptomyces* spores were inoculated in 0.5 ml of 2TY medium, incubated at 50°C for 10 min to induce germination, and then maintained at 30°C for 2–3 h until the donor cells were ready. *E. coli* donor cells were grown in LB medium with kanamycin and apramycin until reaching an OD₆₀₀ of 0.4 - 0.6, then washed three times with cold LB. Five hundred microliters of *Streptomyces* recipient cells were mixed with an equal volume of donor cells and centrifuged to form a pellet. The cell pellet was transferred to a 22-ml SFM plate containing 10 mM MgCl₂. After overnight incubation at 30°C - except for *S. venezuelae*, which was incubated at room temperature – 4.5 ml of SNA medium with nalidixic acid and apramycin was added to the plates, thereafter incubated at 30°C for 5–7 days.

### Experimental identification of Samy *attB* sites in six strains of interest

For each species, three apramycin-resistant exconjugants derived from the conjugation of pOJ260-*SAMYPH94* were isolated and characterized at the genomic level as follows. First, for five strains (*S. ambofaciens* DSM 40697, *S. albidoflavus* J1074, *S. globisporus* NBC_01004, *S. lividans* TK24, and *S. venezuelae* ATCC 10712), one clone per strain was subjected to high-throughput sequencing (PlasmidSaurus) to confirm the unique insertion of the vector (**Data S1**). Next, the insertion identity in three independent clones per strain—covering the five species listed above plus *S. coelicolor* A3(2)—was verified by PCR on genomic DNA (gDNA). The *att*L site was amplified using an SBM585 primer and a species-specific primer, while the *att*R site was amplified using an SBM584 primer and a species-specific primer (**Table S2**). PCR products were analyzed on a 1% agarose gel with ClearSight DNA Stain (Bioatlas). The Thermo Scientific GeneRuler 1 kb Plus DNA Ladder was used as the molecular size marker.

### Analysis of Samy-based vector excision

Three independent clones carrying an integrated form of pOJ260-*SAMYPH94* and three independent stocks of the *S. ambofaciens* ATCC 23877 strain were grown for three days in TSB medium before being collected for phenol extraction of gDNA. To prevent amplification bias due to DNA topology (Hou et al. 2010), both gDNAs and the pOJ260_*SAMYPH94* plasmid were linearized using the SalI restriction enzyme (NEB). Amplification was performed on 5 ng of digested gDNA in a final volume of 10 µl using the LightCycler® 480 SYBR Green I Master (Roche Diagnostics). The primers used to amplify Samy-*att*P junction and *SAMYPH94* gene are listed in **Table S2**. qPCR efficiencies and the absolute number of amplified copies were determined using a standard curve ranging from 3.10^2^ to 3.10^6^ copies of pOJ260_*SAMYPH94* plasmid. The proportion of the plasmid in its excised form was calculated by dividing the *att*P junction copy number by the *SAMYPH94* gene copy number.

### Promoter prediction

We used PromPredict (Rangannan and Bansal 2009) (PromPredict_genome_V1.exe) and G4PromFinder-v.2.1 (Di Salvo 2017; Di Salvo et al. 2018) to predict promoters within the sequence cloned into pOJ260-*SAMYPH94* and pOSV876 vectors, as well as in the region located upstream *SAMYPH94* following its integration at the *att*B site in the *S. ambofaciens* ATCC 23877 genome (GCF_001267885.1). PromPredict_genome_V1.exe was run with default parameter and a window of 50 bp. Please note that *SAMYPH_94* (OR263580.1: 56925-58457) is also referred as *SAM23877_RS39280* under its prophage form within the genome of *S. ambofaciens* ATCC 2387(NZ_CP012382.1: 6589832-6591364).

### Bioinformatic analysis of Samy target gene

The Samy-*att*B logo was generated using the online tool WebLogo (https://weblogo.berkeley.edu/logo.cgi) (Crooks et al. 2004). The gene persistence index was retrieved from Lorenzi *et al*. publication (Lorenzi et al. 2022). Gene synteny between bacterial genomes was analyzed using SyntTax (https://archaea.i2bc.paris-saclay.fr/synttax/) (Oberto 2013) against 19 strains listed in **Fig.2A**. The percentages of identity and similarity between SAM40697_5543 from *S. ambofaciens* DSM 40697 (WP_063483567.1) and SlpB from *B. subtilis* strain 168 (A0A6M3ZAI9) were determined by the Stretcher program, available online (https://www.ebi.ac.uk/jdispatcher/psa/emboss_stretcher) with default parameters. The protein structures were predicted using AlphaFold3 (Abramson et al. 2024). PyMOL (v.2.6.0a0) was used to align the structures and determined the root-mean-square deviation of atomic positions (RMSD). The prediction of *att*B sites was carried out in a panel of 128 genomes corresponding to a panel of 126 genomes representative of the diversity of the genus previously described (Lorenzi et al. 2022), plus 2 strains of interest (*S. lividans* TK24, *S. globisporus* NBC_01004). We used BlastN (ncbi-blast-2.14.0+) to identify sequences similar to the 10 attB sites listed in **Table S3**, as well as the four Samy-attB sites experimentally identified in *S. albidoflavus* J1074, *S. globisporus* NBC_01004, *S. lividans* TK24, and *S. venezuelae* ATCC 10712 (**Fig.1C**). For further analyses, only results with an e-value < 8.10^-4^ and a coverage of at least 70 % were taken into consideration. The origin of replication was identified using Ori-Finder 2022 (Dong et al. 2022)and used to calculate the distance between each putative *att*B site and *ori*C. For each genome, the central compartment is defined as the region between the distal ribosomal operons, while the terminal compartments correspond to the genome’s extremities beyond these operons, as previously described (Lioy et al. 2021; Lorenzi et al. 2022). Data were analyzed with R software (R Core Team 2023) to generate most graphs and perform our statistical tests.

**Figure 1:**
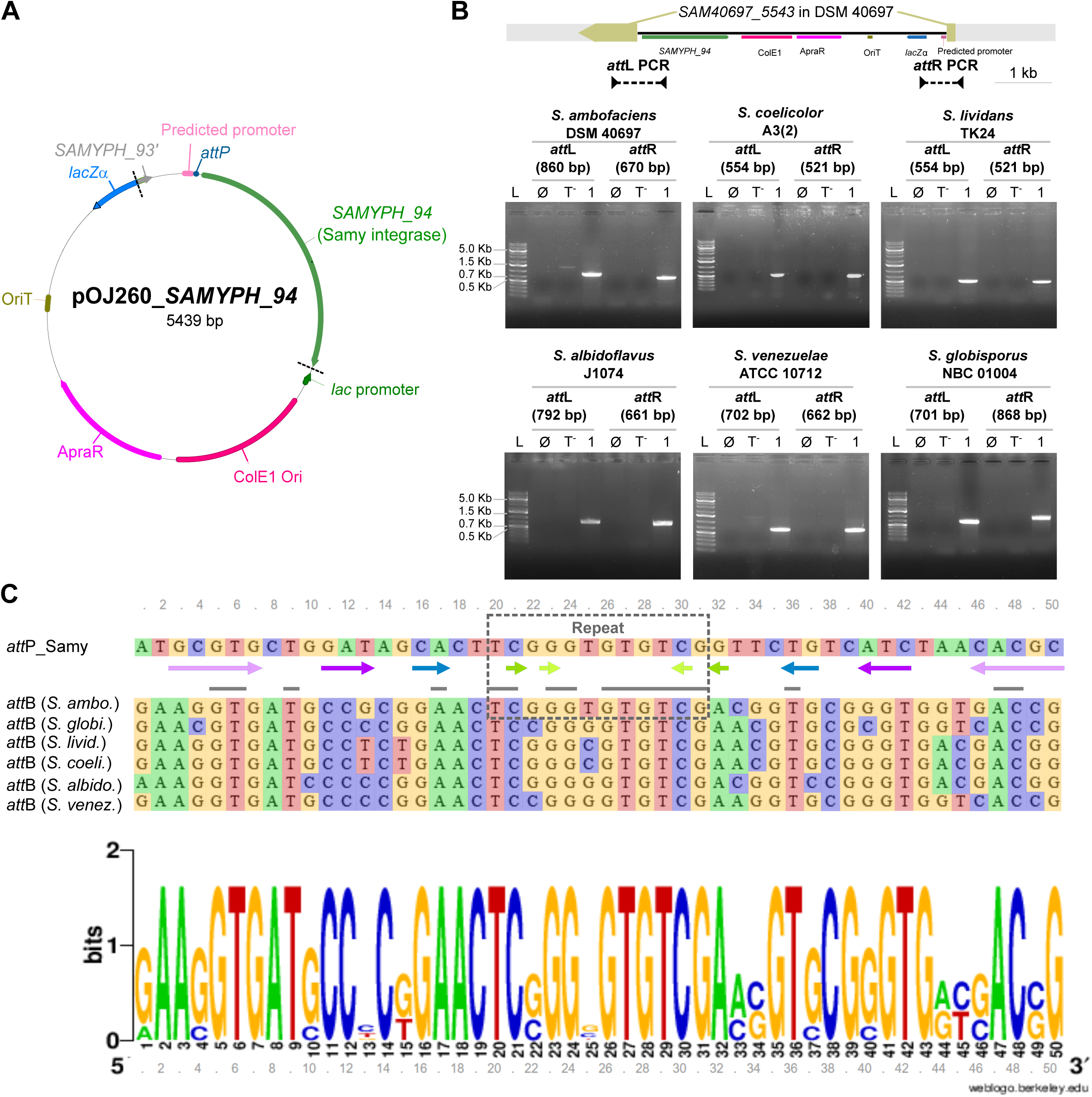
Characterization of Samy phage attachment and integration sites in a panel of six *Streptomyces* species. A. **Schematic representation of pOJ260_*SAMYPH_94* vector.** The two black dashed lines mark the boundaries of the Samy phage region, which was cloned into the pOJ260 vector (Bierman et al. 1992) using a Golden Gate Assembly approach. The predicted promoter and *attP* site sequences are presented in **Fig. S1**. Abbreviations: ApraR (apramycin resistance), ColE1 Ori (ColE1 origin of replication in *E. coli*), *lacZα* (gene encoding the LacZ alpha peptide), OriT (origin of transfer), *SAMYPH_93’* (fragment of *SAMYPH_93* gene). B. **Integration of pOJ260_*SAMYPH_94* into the *SAM40697_5543* gene or its orthologs in a panel of six *Streptomyces* species**. PCR amplification of the *att*L and *att*R regions defined based on Samy prophage orientation in its native *S. ambofaciens* ATCC 23877 host are shown, with expected sizes indicated in brackets. The integration of pOJ260_*SAMPHY_94* in *S. ambofaciens* DSM 40697 is represented at scale. No PCR amplification is expected when gDNA from wild-type *Streptomyces* strains is used as the template (’T-’) or in absence of template (’∅’). The number ‘1’ represents clone #1 analyzed for each strain. ‘L’ denotes the molecular weight ladder (Thermo Scientific GeneRuler DNA Ladder 1Kb Plus). The results for three independent clones per strain is presented in **Fig. S2**. C. **Alignment of Samy-*att*P and *att*B sites from six *Streptomyces* strains against the consensus *att*B sequence**. Sequences of 50 bp centered on the repeat (black hatched rectangle) identified in the integrated form of the Samy prophage (Jaffal et al. 2023) were analyzed. Strain names are presented in abbreviated form before each sequence, with the full names provided in panel B. The logo was generated using the online tool WebLogo (https://weblogo.berkeley.edu/logo.cgi) (Crooks et al. 2004). Gray lines indicate positions that are perfectly conserved across all analyzed sequences. The *att*P site is surrounded by imperfect inverted repeat sequences shown as arrows of the same color, indicating complementary sequences. During integration, a minimal sequence of 2-bp identical between *att*P and *att*B sites is referred to as the crossover sequence (Smith 2015). The repeat region contains several identical 2 bp positions, leaving the exact cleavage site yet to be determined. Please note that *S. coelicolor* and *S. lividans att*B sites are identical.

**Figure 2:**
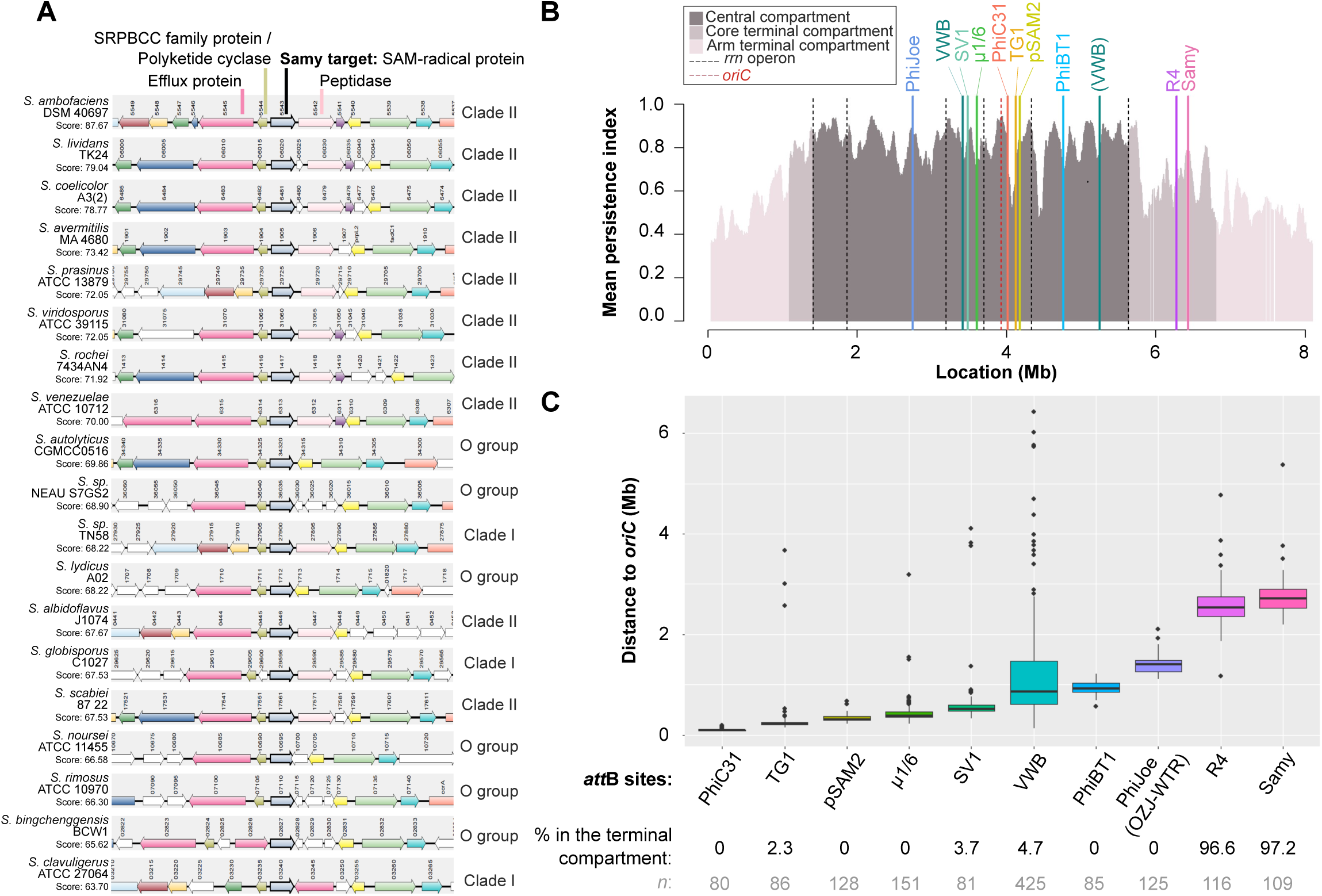
Comparative genomics for the analysis of Samy *att*B sites in a panel of *Streptomyces*. A. **Analysis of gene synteny around *lsr2A* and *lsr2B* genes in a panel of 20 *Streptomyces* strains from Clade I, Clade II and the ‘O’ group.** The genes corresponding to the SAM40697_5543 query protein and its orthologs are drawn in bold. The color code allows the identification of orthologs and paralogs in the different strains. Build using SyntTax (https://archaea.i2bc.paris-saclay.fr/synttax/) (Oberto 2013). *SAM40697_5545* and *SAM40697_5544*, located upstream of *SAM40697_5543*, encode a major facilitator superfamily (MFS) transporter related to multidrug resistance and a polyketide cyclase from the SRPBCC family, respectively. Downstream, *SAM40697_5542* encodes a putative secreted peptidase. **B. Location along *S. ambofaciens* DSM 40697 chromosome of the *attB* sites for all *att-int* systems described to date for engineering the *Streptomyces* chromosome.** The level of gene persistence along the chromosome of *S. ambofaciens* DSM 40697 obtained from (Lorenzi et al. 2022) is represented using a sliding window (81 coding sequences (CDSs), with 1 CDS steps). The positions of all *rrn* operons and of the origin of replication are indicated by dashed black and red lines, respectively. The central compartment (dark gray) is delimitated by the distal *rrn*. Terminal compartments are highlighted in dark pink when they include core genome genes, and in light pink when they do not. Sites that deviate from the consensus sequence (coverage < 80%) are shown in brackets. **Figure S4** presents the same data for four additional strains of interest. C. **Distance of *att*B sites from the origin of replication across 128 *Streptomyces* strains.** The positions of the sites in a panel of 128 strains were determined using BlastN, with the Samy attB sequences identified in this study and all characterized sites from *Streptomyces att-int* systems. Using the OriFinder software to identify the origin of replication, the distance from the origin to each of these sites was calculated. All the boxplots represent the first quartile, median and third quartile. The upper whisker extends from the hinge to the largest value no further than 1.5 * the inter-quartile range (IQR, i.e. distance between the first and third quartiles). The lower whisker extends from the hinge to the smallest value at most 1.5 * IQR of the hinge. The number of sites included in the analysis is displayed below each boxplot, along with the percentage of these sites situated in the terminal compartment.

## Results

### Characterization of the Samy integrase target *att*B sites in a panel of *Streptomyces* species

To evaluate the efficiency and tropism of Samy-based integration system, we cloned, into the conjugative suicide vector pOJ260 (Bierman et al. 1992), the gene *SAMYPH_94* encoding the Samy integrase (510 aminoacids) and its upstream intergenic region (**Fig. 1A**). This latter includes a putative promoter identified using PromPredict software and the ‘TCGGGTGTGTCG’ sequence of the predicted *att*P site (Jaffal et al. 2023) (**Fig. S1.A**). The resulting pOJ260_*SAMYPH_94* was tested for chromosomal integration in six *Streptomyces* hosts representative of the genus’s diversity. These included strains from the Clade II (*Streptomyces albidoflavus* J1074, *S. ambofaciens* DSM 40697, *S. coelicolor* A3(2), *Streptomyces lividans* TK24) and the Clade I (*Streptomyces globisporus* NBC_01004, *Streptomyces venezuelae* ATCC 10712) according to MacDonald and Currie classification (McDonald and Currie 2017; Lorenzi et al. 2022) (**Table S1**). Of note, with an average nucleotide identity of 99.04 % (Lorenzi et al. 2022), *S. ambofaciens* DSM 40697 is closely related to the Samy native host, *S. ambofaciens* ATCC 23877, but devoid of this prophage.

After intergeneric conjugation, the gDNA of one clone from five strains (*S. ambofaciens* DSM 40697, *S. albidoflavus* J1074, *S. globisporus* NBC_01004, *S. lividans* TK24, and *S. venezuelae* ATCC 10712) was subjected to high-throughput sequencing to determine the number and location of the pOJ260_*SAMYPH94* insertion sites. In all cases, the plasmid was integrated in a single copy within the gene orthologous to *SAM40697_5543* (**Table 1, Data S1**) which corresponds to the native location of the complete Samy prophage in *S. ambofaciens* ATCC 23877 (Jaffal et al. 2023). Analysis of the neo-integration site thus allowed to formally determine *att*P, *att*B, *att*L and *att*R sequences (**Table 2**). The uniform definition of the left and right attachment sites for all strains was based on the native orientation of the Samy prophage in its host chromosome. The position of the integration site was confirmed by PCR amplification of the predicted *att*L and *att*R regions in three independent conjugants for each strain - covering the five species listed above plus *S. coelicolor* A3(2) (**Fig. 1B**, **Fig. S2**) - enabling the definition of an *att*B site logo (**Fig. 1C**). These results indicate that the Samy-based integrative vector is functional across a diverse range of phylogenetically distant strains, accommodating variations in the *att*B site.

**Table 1:** Location of Samy-integrase target gene in the panel of six *Streptomyces* strains of interest.

| Strain | Gene name | Chromosomal location* |
| --- | --- | --- |
| *S. albidoflavus* J1074 | XNR_0446 | CP004370.1<br>531548-532585 bp |
| *S. ambofaciens* DSM 40697 | SAM40697_5543 | NZ_CP012949<br>comp(6432936-6434015 bp) |
| *S. coelicolor* A3(2) | SCO6481 | AL645882.2<br>comp(7170631-7171677 bp) |
| *S. globisporus* NBC_01004 | OG449_04345 | CP109059.1<br>999844-1000887 |
| *S. lividans* TK24 | SLIV_06020 | NZ_CP009124.1<br>1344278-1345324 bp |
| *S. venezuelae* ATCC 10712 | SVEN_6313 | NZ_CP029197<br>comp(6889333-6890364 bp) |

\* The start and end of genes containing the Samy-*attB* site are indicated below the chromosome identifier of each strain. 'comp' indicate that the gene is in antisense/complementary orientation.

**Table 2:**
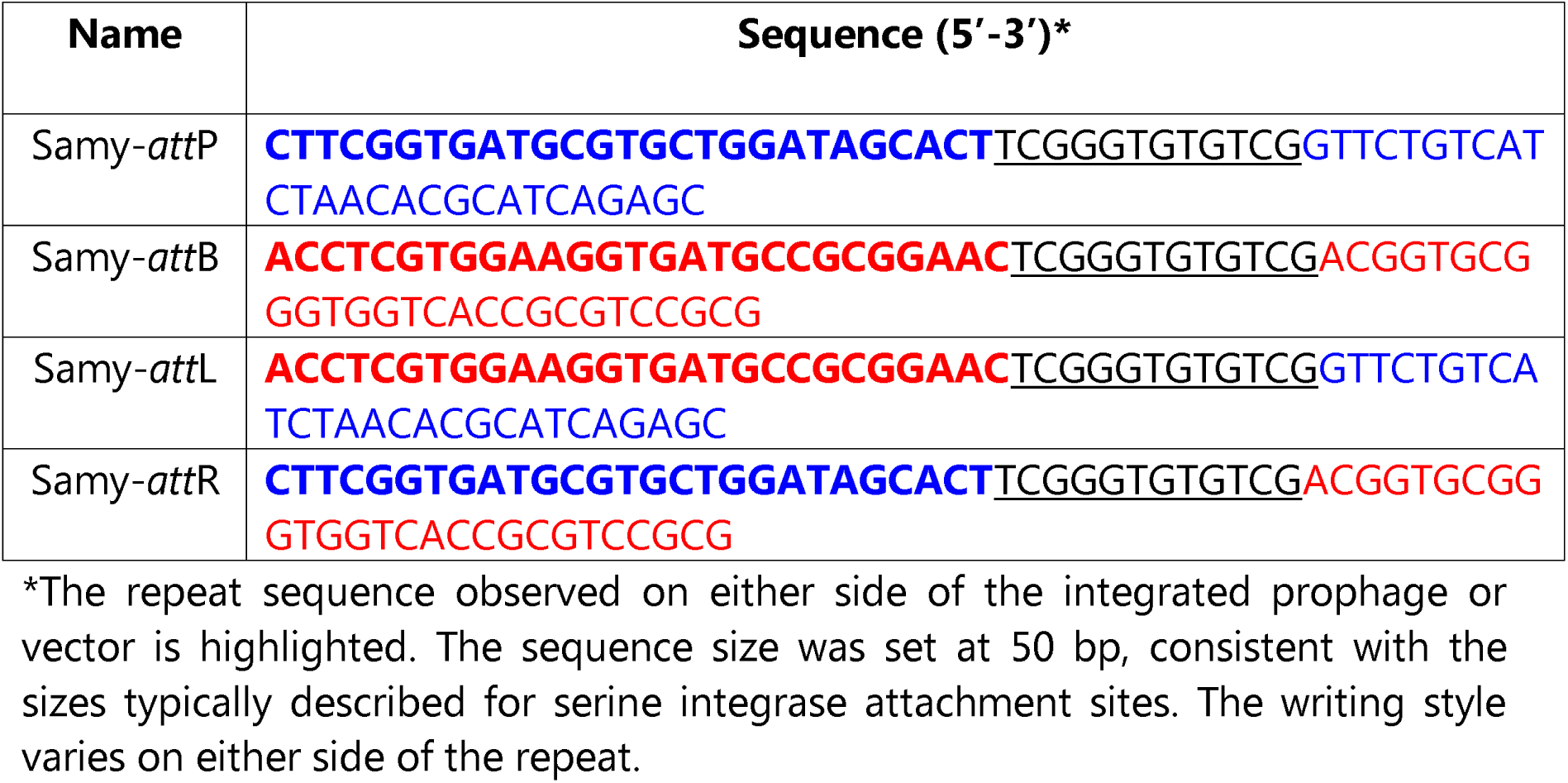
Sequences of the Samy integrase attachment sites in the phage and in the *S. ambofaciens* chromosome.

### Bioinformatic analysis of the Samy-*att*B integration site

The genes targeted by Samy integrase, *SAM40697_5543* and its orthologs, encode a radical S-adenosyl-L-methionine (SAM) protein belonging to the Rv2578c family. This iron-sulfur cluster binding protein is annotated as a putative SplB (spore photoproduct lyase B) in several strains. The SlpB protein is involved in DNA repair in *Bacillus subtilis* (Slieman et al. 2000). The similarity with this protein is rather low: SAM40697_5543 and the *B. subtilis* SlpB protein share 17.6% identity and 29.2% similarity over 387 amino acids, with 18.9% gaps. Since proteins with sequence identity below 25% can still have similar structures, we compared the 3D structures of SAM40697_5543 and SlpB from *B. subtilis* using AlphaFold3 (**Fig. S3)**. Given that the root mean square deviation (RMSD) between these structures exceeds 4 Å, the annotation of Samy’s target gene as SlpB should be interpreted with caution and requires experimental validation.

The persistence of *SAM40697_5543* gene in a previously characterized panel of 127 *Streptomyces* strains is around 88% (Lorenzi et al. 2022). Furthermore, we observed that the synteny around this gene is relatively well conserved in a panel of twenty strains chosen in Clade I, Clade II and group ‘O’ (others) to represent the *Streptomyces* genus diversity (McDonald and Currie 2017; Lorenzi et al. 2022) (**Fig.2A**). *SAM40697_5543* orthologs are frequently co-conserved with their upstream divergent gene, which encodes a putative polyketide cyclase. In certain strains such as *S. ambofaciens* DSM 40697, the target gene of the Samy integrase could possibly be expressed in operon with genes encoding a peptidase (**Fig.2A**). In this case, the integration of Samy might have a polar effect on the expression of the downstream gene.

Finally, we conducted a BlastN analysis to determine the localization of Samy-*att*B in a panel of 128 genomes representative of the genus diversity. Using the *att*B sites identified in our strains of interest as input (**Fig.1C**), Samy-*att*B sites were predicted in 83% of the strains, as a single copy (**Table S4**). In parallel, we have also analyzed the positions within these genomes of all the other integration systems described for genetically modifying *Streptomyces* (**Table S3**). This analysis reveals that the Samy-*att*B site is located in the terminal compartment of most of the analyzed strains (**Table S4**), with only a few exceptions (*Streptomyces bingchenggensis* BCW 1, *Streptomyces fungicidicus* TXX3120, *Streptomyces sp.* 11 1 2) whose genome was previously described as highly recombined (Lorenzi et al. 2022). Comparison to other sites *att*B sites (**Table S3 & S4**) reveals that the *att*B-Samy site is located furthest from the origin of replication in most strains (**Fig. 2B & C, Fig S4**).

Taken together, these results indicate that the Samy-*att*B site within a gene encoding a SAM-radical protein is frequently present across the genus, as a single copy and located in the terminal compartment of the *Streptomyces* chromosome.

### Comparative analysis of the integration via PhiC31- and Samy-based vectors

The closest known phage to Samy is PhiC31 (Jaffal et al. 2023). However, while Samy integrates within the terminal compartment, PhiC31 typically integrates near the origin of replication in most species (**Fig. 2B**). To compare the efficiency of integration-vectors based on both types of integrases, we introduced the Samy *att*P and integrase encoding gene between SbfI and AflII sites in the pOSV backbone of the plasmid collection built by Aubry and colleagues (Aubry et al. 2019). This allows for the generation of perfectly comparable plasmid vectors, pOSV802 (PhiC31-based) and pOSV876 (Samy-based), that differ only in their integrative modules (**Fig. 3A**). These vectors were introduced through intergeneric conjugation into the *S. albidoflavus* J1074 and *S. lividans* TK24 strains. The Samy-*att*B sites in these two strains each exhibit 84% identity to the native *att*B sequence in *S. ambofaciens*, though they differ at distinct positions. The efficiency of conjugation by the pOSV876 vector was equivalent – around 10^3^ exconjugants per colony forming unit (CFU) - between these two host strains, whereas the transduction efficiency of pOSV802 varies by several orders of magnitude between them. Indeed, the pOSV876 conjugation efficiency was one-log higher than that of pOSV802 vector when the recipient bacteria were *S. albidoflavus* J1074, but lower than that of pOSV802 vector when the recipient was *S. lividans* TK24 (**Fig. 3B & 3C**). This suggests that host-specific factors influence the efficiency of PhiC31 integrative vector-based gene transfer. In particular, the PhiC31-*att*B site in *S. albidoflavus* J1074 shows 91.4% identity to the reference sequence, whereas it is perfectly identical in the *S. lividans* TK24 chromosome (**Table S4**). This difference may account for all or part of the difference in efficiency between the strains.

**Figure 3:**
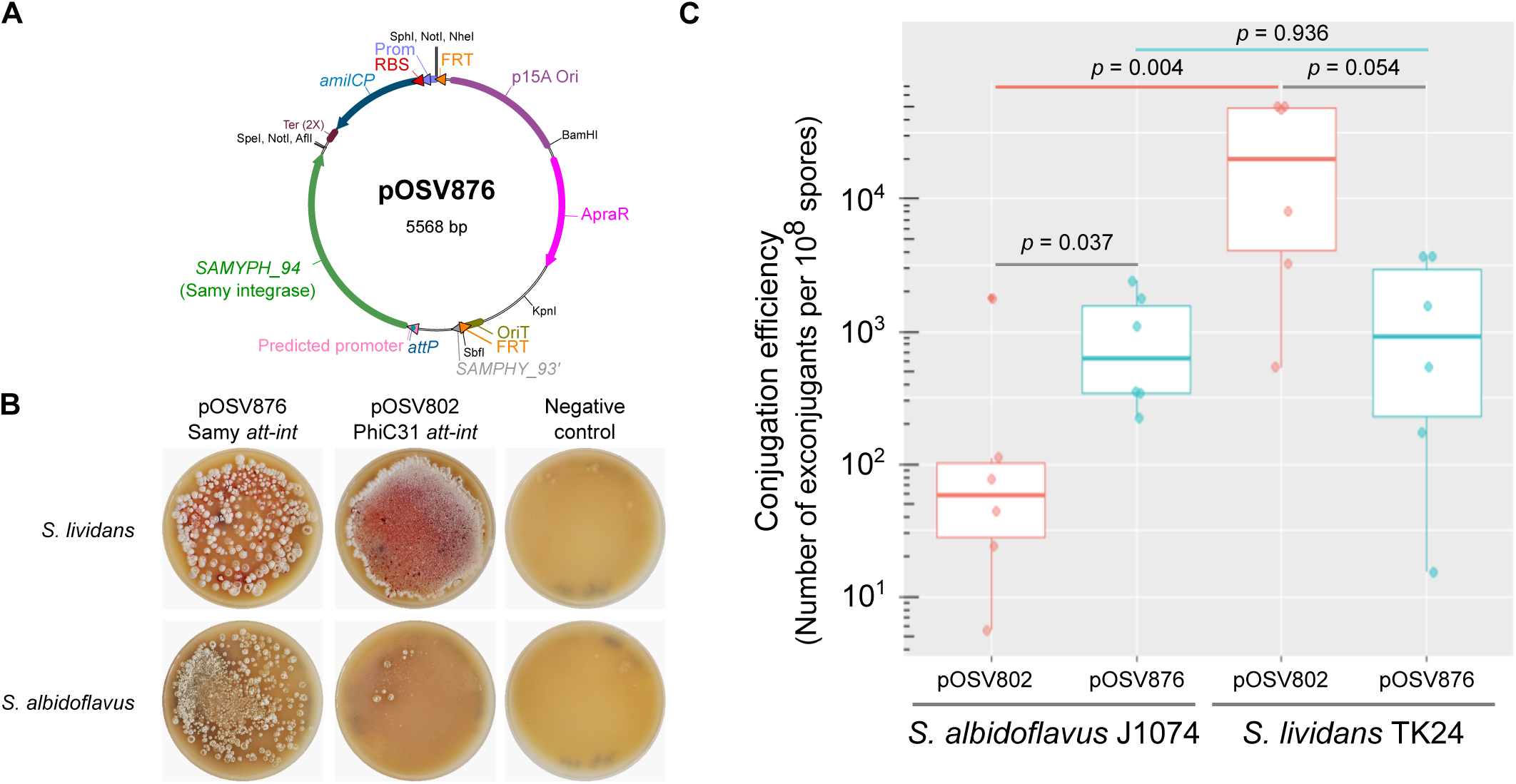
Comparative analysis of Samy- and PhiC31-based integrative system. **A. Schematic representation of pOSV876 vector.** The vector is organized in a modular fashion according to the collection described by Aubry and colleagues (Aubry et al. 2019). The predicted promoter and *attP* site sequences are presented in **Fig. S1**. Abbreviations: *amilCP* (gene encoding an *Acropora millepora* blue chromoprotein), ApraR (apramycin resistance), FRT (flippase recognition target), OriT (origin of transfer), p15A Ori (p15A origin of replication in *E. coli*), Prom (*E. coli* promoter), RBS (ribosome binding site), *SAMYPH_93’* (fragment of *SAMYPH_93* gene), Ter (terminators). **B. A representative result of a conjugation experiment of *S. albidoflavus* J1074 and *S. lividans* TK24 using an *E. coli* strain to transfer pOSV 873 and pOSV802**. In this latter plasmid, PhiC31 *att-int* system replaces Samy system (Aubry et al. 2019). An *E. coli* strain devoid of conjugative plasmid was used as a control. The plates show the exconjugants obtained by spreading a 100-fold dilution from a conjugation experiment conducted with approximately 10^8^ spores. **C. Conjugation efficiency of an integrating vector, containing Samy or PhiC31 *att-int* systems into *S. albidoflavus* J1074 and *S. lividans* TK24.** Boxplots are represented as in **Fig2C**. Each dot represents an independent experiment. The *p* values of two-sided Wilcoxon rank sum tests with continuity correction are indicated.

Overall, these results suggest that the efficiency of integration and/or expression of the resistance cassette in the terminal compartment using the Samy-based vector is relatively high, comparable to the PhiC31 integrative system which varies by strain.

### Samy integrase-based vector exhibits a basal level of excision *in vivo*

To assess whether Samy integrase-based integration is unidirectional, we quantified the proportion of *att*P junctions in apramycin-resistant exconjugants following the introduction of pOJ260*_SAMYPH_94*. Interestingly, the number of *att*P junction per genome varied between approximately 10⁻⁶ and 10⁻³, depending on the strain, with *S. coelicolor* exhibiting the highest levels of excision (**Fig. 4**). We further examined this activity within the native context of the Samy prophage in the *S. ambofaciens* ATCC 23877 strain. Notably, we observed a low level of spontaneous excision, comparable to that of pOJ260_*SAMYPH_94* in *S. ambofaciens* DSM 40697 strain, suggesting that the excision activity remains consistent and is not significantly influenced by the presence of other prophage genes.

**Figure 4:**
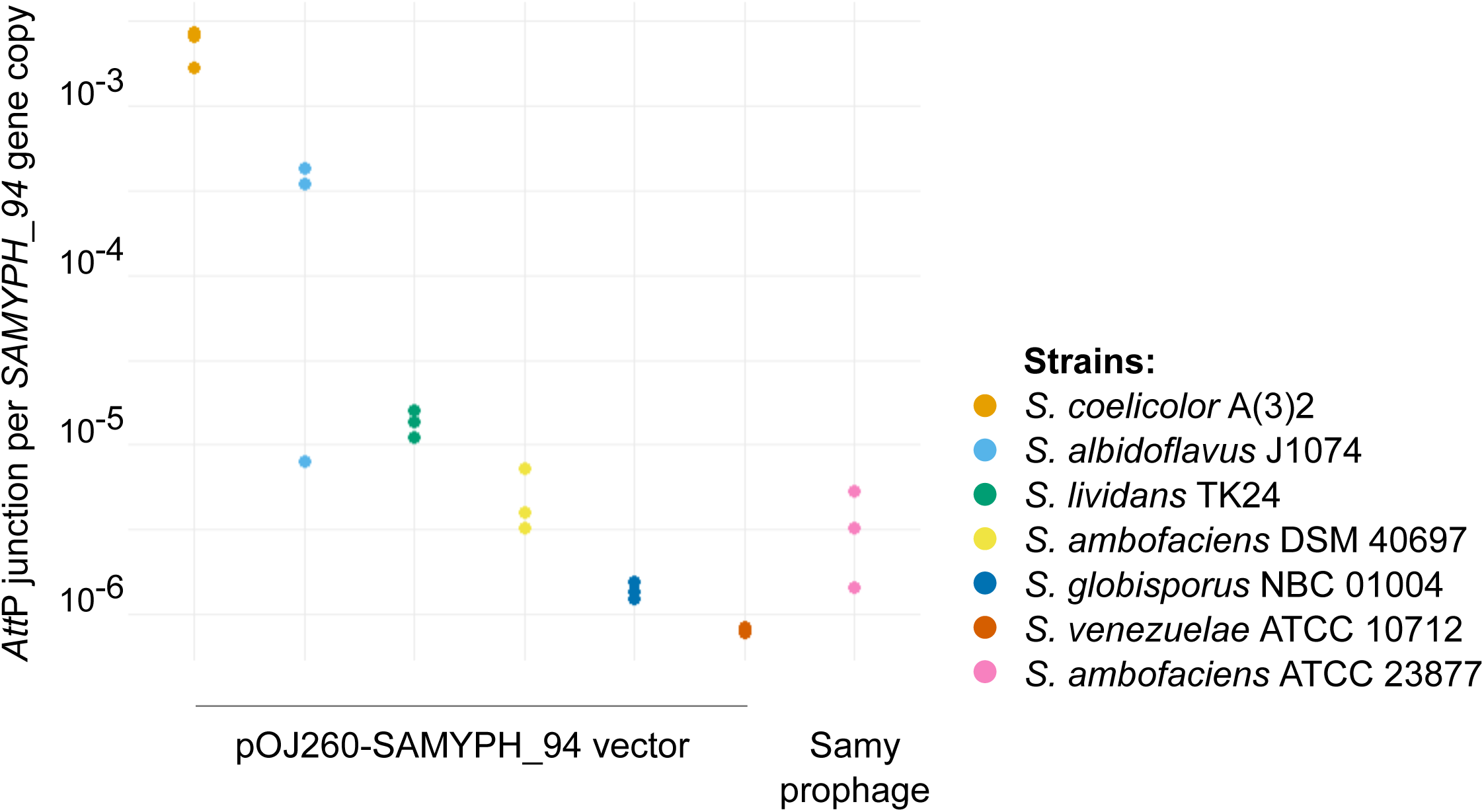
Basal excision level of Samy integrase-based vector and native prophage. The excision levels of the pOJ260_*SAMYPH_94* vector and the Samy prophage were assessed by quantitative PCR (qPCR) performed on gDNA from six *Streptomyces* strains in which the pOJ260-*SAMYPH_94* vector was integrated, as well as from the native strain containing the complete Samy prophage. For each condition, the *att*P junction copy number was quantified relative to the *SAMYPH_94* gene copy number. The results from three quantifications performed on distinct clones are presented.

Upon integration, the *SAMPHY94* gene becomes separated from its native viral promoter – located just upstream the *att* repeat (**Fig 1B, Fig S1A**). However, on the *att*L side, promoters upstream of the *SAMYPH_94* gene have been predicted (**Fig S1B**). This finding aligns with previous studies analyzing the Samy transcriptome, which identified *SAMPHY_94* as one of the genes consistently expressed across 13 different culture conditions and growth stages (Jaffal et al. 2023).

Altogether, these results suggest that the Samy integrase remains expressed even after integration and may exhibit a basal excisionase activity, influenced by the genetic background and/or host factors. Such an activity has been previously described for other serine-recombinases as PhiJoe (Fogg et al. 2017) and PhiBT1 (Zhang et al. 2008).

## Discussion

In this study, Samy integration was experimentally validated in six strains and explored using BLAST prediction in a panel of 128 strains. The latter approach has limitations, as some vectors may integrate at pseudo-*att*B sites in the absence of the cognate *att*B site (Combes et al. 2002; Baltz 2012). This capability has been used, for instance, to integrate PhiC31-based tools at the pseudo-*attP* sites in the human genome (Thyagarajan et al. 2001). Conversely, the presence of the *att*B site does not necessarily guaranty an efficient integration, as described for SV1- (Fayed et al. 2014) or PhiWTR- (Ko et al. 2020) based integrative vectors in some strains. The factors involved in the restriction of target cell tropism remain to be elucidated in these cases. Within the above-mentioned limits, our bioinformatics analysis indicates that the Samy integrase-based tools enable integration furthest from the origin of replication compared to other integration tools (**Fig. 2**). Consequently, they extend the range of chromosome regions that can be potentially targeted in *Streptomyces*.

The ideal genetic tool integrates into a “neutral” site, particularly in terms of functions related to fitness and metabolic capacity in *Streptomyces*. This is illustrated by the few tools that restore post-integration the functionality of the target-encoding gene [*e.g*., µ1/6 (Farkašovská and Godány 2012), pSAM2 (Boccard et al. 1989)]. However, in most cases the target gene is inactivated. When this occurs, the impact of integration may not be neutral and requires thorough investigations to be accurately characterized. For instance, integration mediated by PhiC31 has been shown to disrupt beta-oxidation metabolism in *S. ambofaciens* ATCC 23877 (Talà et al. 2018). Similarly, integration at the *att*B-PhiBT1 site can cause a polar effect on downstream genes, leading to defects in cell differentiation and delayed spore germination (Gonzalez-Quiñonez et al. 2016). Therefore, dedicated experimental studies are crucial to assess the consequences of integration. Samy target gene, *SAM40697_5543*, encodes a SAM radical protein, akin to PhiJoe, PhiOZJ and PhiWTR phages which integrate into a distinct SAM radical protein encoding gene (Ko et al. 2020). In some strains, the function of the gene containing *att*B-Samy is tentatively annotated as potentially involved in DNA repair, based on sequence similarity with SplB. However, this similarity is not strong (**Fig. S3**), and any connection to DNA repair remains speculative, especially since the surrounding genes at the *att*B site do not appear to be linked to this process. The *SAM40697_5543* gene is inactivated in the environmental strain *Streptomyces ambofaciens* ATCC 23877, which is Samy’s native host strain. This suggests that its inactivation does not significantly impact bacterial fitness in nature. Moreover, another integrative element is located a few hundred base pairs upstream of *att*B-Samy, indicating that this region serves as an integration hotspot.

Integrative vectors based on the Samy integrase, like the native virus, exhibit a basal excision level. Although considered as rare in serine integrases, this bidirectionality has also been reported for PhiJoe (Fogg et al. 2017) and PhiBT1 integrases (Zhang et al. 2008). Fogg and colleagues (Fogg et al. 2017) proposed that the elevated basal excision activity of _Joe integrase may result from partial inhibition of synapsis by the coiled-coil motif when the integrase is bound to *att*L and *att*R, reminiscent of the hyperactive _C31 mutant IntE449 (Rowley et al. 2008). The basal excision level observed with the Samy integrase-derived vector is particularly high in the *S. coelicolor* strain (**Fig. 4**). This observation suggests that host factors—such as partner proteins and/or the expression level of the integrase in its integrated form—may influence this phenomenon. These results highlight the need for a more systematic evaluation of the basal excision level of integrative vectors across multiple *Streptomyces* strains.

Novel integrating vectors are desirable because differences in genome architecture and host-factors between strains can limit the effectiveness of existing vectors in specific actinomycetes. Additionally, expanding the repertoire of genetic tools will encourage combinatorial engineering efforts (Li et al. 2017; Li et al. 2019; Gao and Smith 2021). To complete the suite of *Streptomyces int/attP* vectors, we constructed apramycin-resistant derivatives of pSET152 and pOSV vectors harboring Samy-integrase.

Finally, variations in gene expression depending on their natural chromosomal location (Lioy et al. 2021; Lorenzi et al. 2022) or their integration site (Phelan et al. 2017) further justify the need to enhance the genetic toolbox for *Streptomyces* and other actinomycetes. Naturally, SMBGCs tend to be enriched in the terminal compartments of *Streptomyces* chromosomes (Lioy et al. 2021; Lorenzi et al. 2022). The impact of this localization on their cryptic nature and the strength of their repression or activation in response to environmental conditions remains to be explored. Currently, tools based on Samy and R4 integrases are the only ones targeting the terminal compartment in most strains. Therefore, they are valuable for investigating these questions.

Similarly to pSAM2, the first exploited integrative tool described in *Streptomyces*, (Boccard et al. 1989; Raynal et al. 1998), our study demonstrates that integrative systems can be discovered through an approach guided by the study of integrated genetic elements such as prophages. This complements the more traditional method of isolating phages from the environment. Hundreds of complete or defective prophages have been identified in *Streptomyces* genomes (Sharma et al. 2023), highlighting their potential as a valuable source of genetic tools. Exploiting these elements requires the amplification and cloning of their integrases and putative *attP* sites, as predicted by identifying the repeats bordering the prophages. To achieve this, the conditions that activate the excision of these integrated mobile genetic elements have to be identified, as we did earlier for the Samy prophage. However, this may constitute a bottleneck for exploiting these systems. Alternatively, artificial excision can be used to obtain circular prophage form. This required for the simultaneously cloning the integrase and the predicted excision factor. With this approach, the cloning of the excision tool is a prerequisite for exploiting the integrative potential of prophages and integrative mobile genetic elements. Finally, synthetic biology provides the option to directly synthesize genes encoding integrases and their predicted *att*P sites. Collectively, these different strategies offer options for expanding the toolbox for precisely integrating and excising of (over)cassettes into genomes.

## Supporting information

DataS1

## Acknowledgments

We thank Pascaline Tirand for her daily help. We would also like to thank Hervé Leh, Virginia Lioy and Frédéric Boccard for their helpful discussions. While preparing this manuscript, the authors utilized the ChatGPT service to rephrase certain sentences and enhance the overall quality of the English language.

## Declarations

### Clinical trial number

Not applicable.

### Ethical Approval

Not applicable.

### Availability of data and materials

The *att*B sequences used in this study are detailed in **Fig. 1C** and **Table S3**. The bioinformatics analyses supporting some of the figures are summarized in Table S3. The integrative vectors generated in this study are available upon request from the corresponding author (SBM).

### Author Contribution

SBM conceived and designed the research and drafted the manuscript. ND, FRP, and HJ performed the experiments. All authors reviewed and approved the final manuscript. The FASTA files of genome assemblies of exconjugants derived from the conjugation of pOJ260-*SAMYPH94* are available in Data S1.

