## Supplementary information for "A new-engineered integrative tool to target the terminal compartment of the *Streptomyces* chromosome"

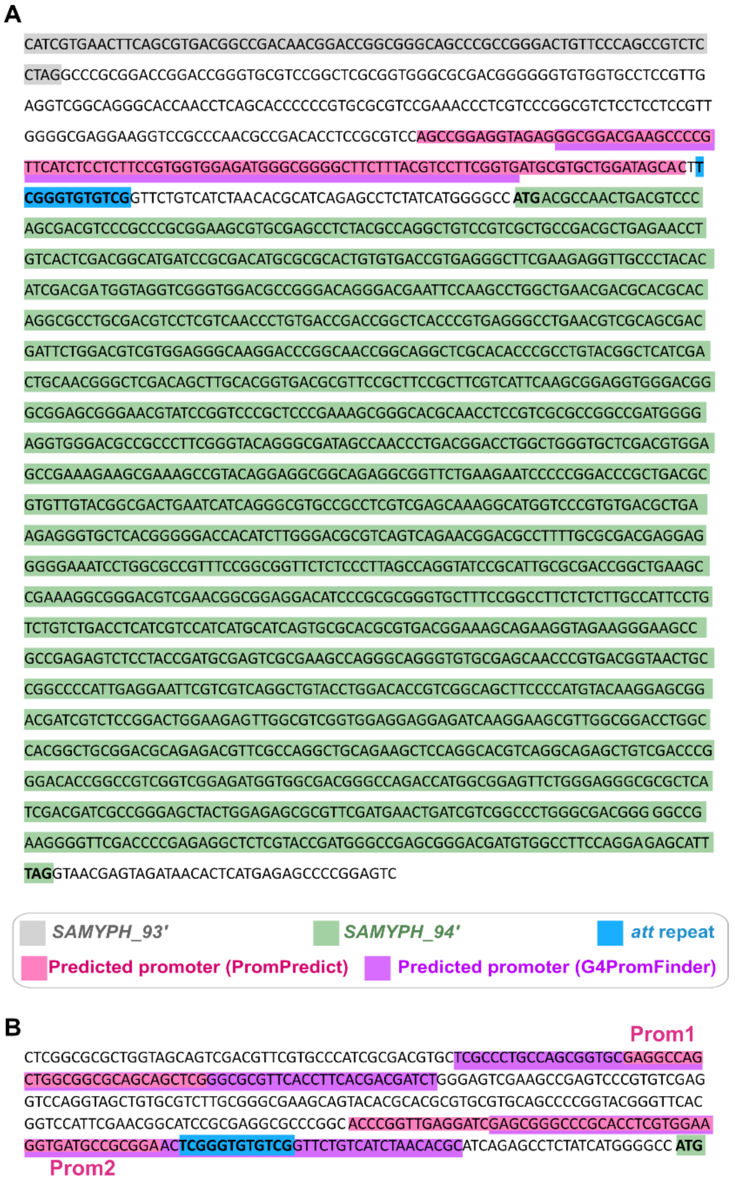


**Figure S1: Samy integrase gene and its upstream sequences**

A. **Sequence cloned in pOJ260_*SAMPHY_94* and pOSV876 vectors.** The promoter predicted with a confidence level of 4 using PromPredict software (version 1, (Rangannan and Bansal 2009)) overlaps with the promoter predicted by G4PromFinder (Di Salvo 2017; Di Salvo et al. 2018). Start and stop codons as well as the repeat present in *att* sites (**Table 2**) are written in bold. The color code is indicated below the sequence.

B. **Sequence upstream *SAMPHY_94* gene after integration at the *S. ambofaciens att*B site.** The region corresponds to the *att*L side shared by the ATCC 23877 and DSM 40697 strains. The promoters 'Prom1' and 'Prom2'O were predicted on the ATCC 23877 strain by the PromPredict software (version 1, (Rangannan and Bansal 2009)) with confidence levels of 2 and 3, respectively. They both overlap with the promoters predicted by G4PromFinder (Di Salvo 2017; Di Salvo et al. 2018). The color coding follows the scheme used in panel A. Only the start codon of *SAMPHY_94* is indicated.

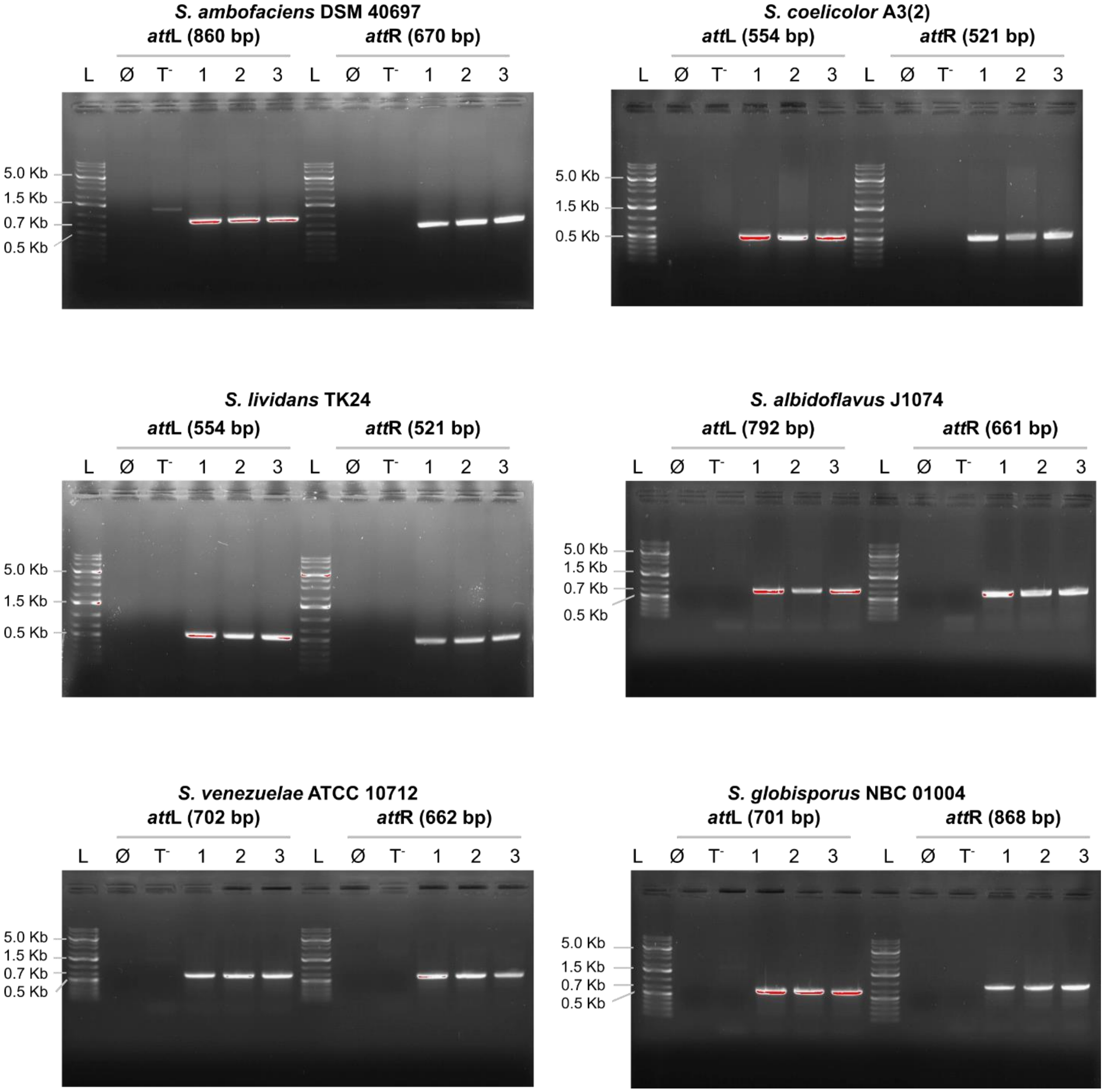


**Figure S2: Analysis of the integration site in three independent clones per strain of interest**

As described in **Fig 1B,** PCR amplification of the *att*L and *att*R regions defined based on Samy prophage orientation in its native *S. ambofaciens* ATCC 23877 host are shown, with expected sizes indicated in brackets. No PCR amplification is expected when genomic DNA from wild-type *Streptomyces* strains is used as the template ('T-') or in absence of template ('∅'). The numbers represent distinct clones analyzed for each strain. 'L' denotes the molecular weight ladder (Thermo Scientific GeneRuler DNA Ladder 1Kb Plus).

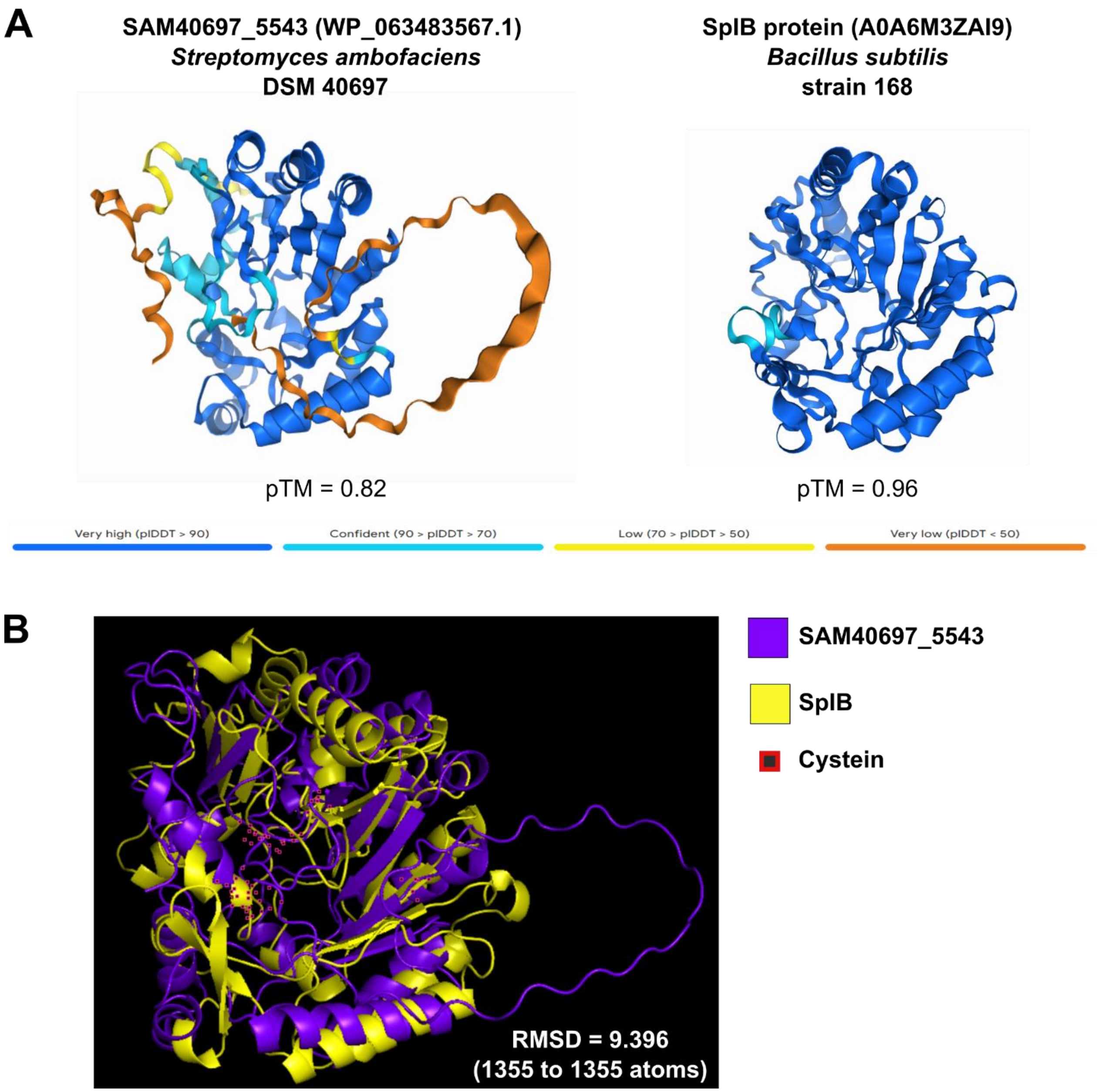


**Figure S3: Comparative analysis of the AlphaFold 3 predicted structures of SAM40697_5543 from *S. ambofaciens* DSM 40697 and SlpB from *B. subtilis* strain 168**

Structures are shown individually in panel A and overlaid in panel B. The RMSD (root-mean-square deviation of atomic positions) was calculated by PyMOL (v.2.6.0a0). The number of positions taken into account for the calculation is indicated in each case. AlphaFold produced a per-residue model confidence score (pLDDT) between 0 and 100. Some regions below 50 pLDDT may be unstructured in isolation.

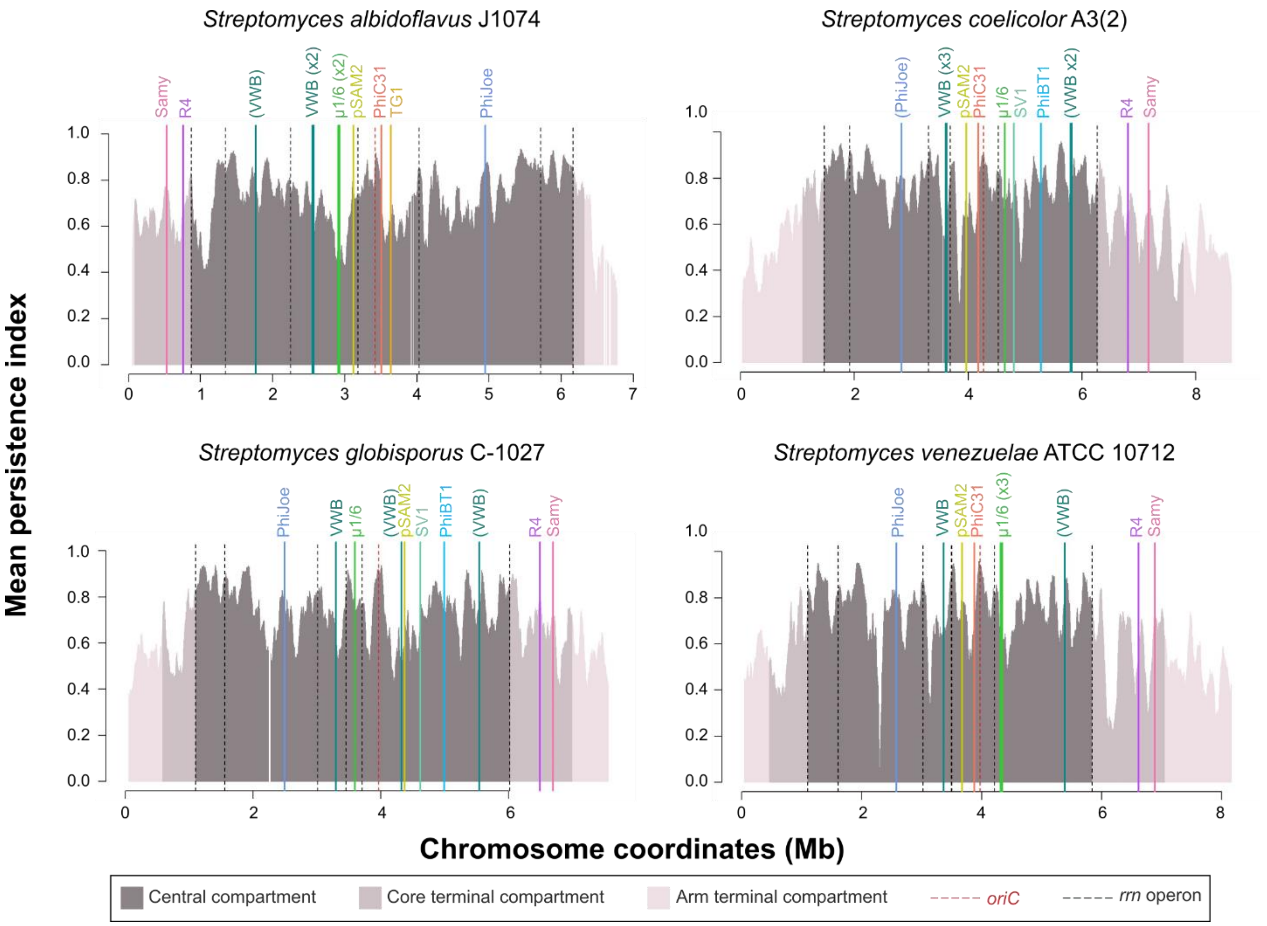


**Figure S4: Chromosomal locations of *attB* sites for all *att-int* systems identified to date for engineering the *Streptomyces* chromosome in four strains of interest**

The level of gene persistence along the chromosome obtained from (Lorenzi et al. 2022) is represented using a sliding window (81 coding sequences (CDSs), with 1 CDS steps). The positions of all *rrn* operons and of the origin of replication are indicated by dashed black and red lines, respectively. The central compartment (dark gray) is delimitated by the distal *rrn*. Terminal compartments are highlighted in dark pink when they include core genome genes, and in light pink when they do not. Sites that deviate from the consensus sequence (coverage < 80%) are shown in brackets. Numbers indicate the frequency of site occurrence.

### Table S1: Strains and plasmids used in this study

| Name | Main characteristics | Reference |
|---|---|---|
| ***Streptomyces* strains*** | | |
| ***S. ambofaciens* ATCC 23877** | RP3486 strain deposited at the ATCC by RP; Sequenced genome: GCF_001267885.1_ASM126788v1 | (Pinnert-Sindico S, Ninet L, Preud'homme J, Cosar C 1954), genome sequence (Thibessard et al. 2015), Pernodet-Lautru's lab collection |
| ***S. ambofaciens* DSM 40697** | Tü13 strain deposited at the DSM collection; Sequenced genome: CP012949 | (Hütter 1967), genome sequence (Thibessard and Leblond 2016), Pernodet-Lautru's lab collection |
| ***S. albidoflavus* J1074** | Sequenced genome: CP004370 | (Zaburannyi et al. 2014), Pernodet-Lautru's lab collection |
| ***S. coelicolor* A(3)2** | Sequenced genome: NC_003888.3 | (Freeman et al. 1977), Pernodet-Lautru's lab collection |
| ***S. globisporus* NBC_01004** | Sequenced genome: CP109059 | Pernodet-Lautru's lab collection |
| ***S. lividans* 66 TK24** | Streptomycin-resistant mutation (*str6)* genetic marker<br>Sequenced genome: CP009124 | (Hopwood et al. 1983), Pernodet-Lautru's lab collection |
| ***S. venezuelae* ATCC 10712** | *S. venezuelae* Ehrlich *et al.*, type strain isolated from soil in Venezuela<br>Sequenced genome: CP029197 | Pernodet-Lautru's lab collection |
| ***Escherichia coli* strain** | | |
| **DH5α** | General cloning strain | Lab collection |
| **ET12567 pUZ8002** | Donor strain, whose conjugation helper non transmissible plasmid (pUZ8002) encodes kanamycin resistance, used for the conjugative transfer of DNA from *E. coli* to *Streptomyces* | (Gust et al. 2004) |
| **Plasmids§** | | |
| **pOJ260** | **ColE1, *oriT, lacZ'*, *aac(3)IV***<br>Non-integrative plasmid, replicative in *E. coli* but not *Streptomyces*, carrying the apramycin resistance gene and an origin of transfer for conjugation | (Bierman et al. 1992) |
| **pOJ260-*SAMYPH94*** | **ColE1, *oriT, lacZ'Ω SAMYPH94*, *aac(3)IV*** | This study |

| | | |
|---|---|---|
| | Samy-based integrative vector, derivative of pOJ260, containing, between the *lac* promoter and *lacZ'* sequence, the 56,510-58,494 genomic region of Samy phage (OR263580.1) including the *SAMYPH94* gene encoding the Samy-integrase, its promoter and Samy-*att*P site sequences | |
| **pOSV802** | **p15A*, oriT, amilCP*, *aac(3)IV, PhiC31 int-attP,* FRT**<br>PhiC31-based integrative vector | (Aubry et al. 2019) |
| **pOSV876** | **p15A*, oriT, amilCP*, *aac(3)IV, Samy int-attP*, FRT**<br>Samy-based integrative vector, new derivative of pOSV vector, containing between *Afl*II and SbfI sites the 56,510-58,494 genomic region of Samy phage (OR263580.1) including the *SAMYPH94* gene encoding the Samy-integrase, its promoter and Samy-*att*P site sequences | This study |

*Abbreviations: ATCC (American Type Culture Collection), DSM (*Deutsche Sammlung von Mikroorganismen*), RP (Rhône-Poulenc)

§ *aac(3)-IV*, apramycin resistance gene; *amilCP*, gene coding for an *Acropora millepora* blue chromoprotein; *att*P, phage attachment site; *ColE1*, origin of replication in *E. coli* (not functional in *Streptomyces*); FRT corresponds to the sites recognized by the Flp Recombinase; *int*, phage integrase gene; *lacZ'*, gene encoding the LacZα; *oriT*, origin of transfer; *p15A*, origin of replication in *E. coli* (not functional in *Streptomyces*)

**Table S2: Primers used in this study**

| Primer name | Target region | Sequence (5′-3′)* |
|---|---|---|
| **Primers used for vector cloning** | | |
| **SBM375** | pOJ260 | TTTGGTCTCCCCTGTGTGAAATTGTTATCC |
| **SBM376** | pOJ260 | CAGCTATGACGGTCTCATGATTACGAATTCGATATCG |
| **SBM560** | Upstream *SAMYPH94* (56,510-56,529 genomic region of Samy phage OR263580.1 sequence) | GGGGGTCTCTAACACATCGTGAACTTCAGCGTGA |
| **SBM561** | Downstream *SAMYPH94* (58,477-58,494 genomic region of Samy phage OR263580.1 sequence) | CGCGGTCTCGCAGGGACTCCGGGGCTCTCATG |
| **SBM588** | Downstream *SAMYPH94* (58,475-58,494 genomic region of Samy phage OR263580.1 sequence) | CCGCTTAAGGACTCCGGGGCTCTCATGAG |
| **SBM589** | Upstream *SAMYPH94* (56,510-56,528 genomic region of Samy phage OR263580.1 sequence) | TTCCCTGCAGGCATCGTGAACTTCAGCGTG |
| **Primers used for Samy-*att*L and Samy-attR region amplifications** | | |
| **SBM584** | Samy-*att*P region on pOJ260-*SAMYPH94*, divergent primer used to amplify *att*R | CTGCAGGTCGACTCTAGAGG |
| **SBM585** | Samy-*att*P region on pOJ260-*SAMYPH94*, divergent primer used to amplify *att*L | CCTCCGCTTGAATGACGAAG |
| **HL15** | *S. ambofaciens* DSM 40697 and ATCC 23877 strain genomic regions located upstream Samy-*att*B site, used to amplify *att*L | GATCAGCGTGCCCTTGGT |
| **HL16** | *S. ambofaciens* DSM 40697 and ATCC 23877 strain genomic regions located downstream Samy-*att*B site, used to amplify *att*R | TCCTCCAGGGCCTGGTTC |

| | | |
|---|---|---|
| **SBM582** | *S. coelicolor* A(3)2 and *S. lividans* TK 24 strain genomic regions located downstream Samy-*att*B site, used to amplify *att*R | CCTCAATGCGCTGGGAGAA |
| **SBM583** | *S. coelicolor* A(3)2 and *S. lividans* TK 24 strain genomic regions located upstream Samy-*att*B site, used to amplify *att*L | CACCCGGTTCAGGATCGAG |
| **SBM590** | *S. albidoflavus* J1074 genomic region located downstream Samy-*att*B site, used to amplify *att*R | ATGTCCACTTCCTCCAGCAC |
| **SBM591** | *S. albidoflavus* J1074 genomic region located upstream Samy-*att*B site, used to amplify *att*L | GGTAGCAGTTGACGTTGGTG |
| **SBM592** | *S. globisporus* NBC_01004 genomic region located downstream Samy-*att*B site, used to amplify *att*R | ATGTGGAGCCGGTTCTTGAG |
| **SBM593** | *S. globisporus* NBC_01004 genomic region located upstream Samy-*att*B site, used to amplify *att*L | CTTGACGACGATCTGGGAGT |
| **SBM594** | *S. venezuelae* ATCC 10712 genomic region located downstream Samy-*att*B site, used to amplify *att*R | ATTGACTTCCTCCAGCTCGT |
| **SBM595** | *S. venezuelae* ATCC 10712 genomic region located downstream Samy-*att*B site, used to amplify *att*R | TCTTGACGACGATCTGGGAG |
| **Primers used for qPCR analyses** | | |
| **SBM609** | Samy-*att*P junction | CGGGGCTTCTTTACGTCCTT |
| **SBM615** | Samy-*att*P junction | GTAGAGGCTCGCACGCTT |
| **SBM611** | *SAMYPH94* (Samy integrase) | GACGTCCTCGTCAACCCTG |
| **SBM612** | *SAMYPH94* (Samy integrase) | ACCTCCGCTTGAATGACGAA |

* Restriction sites are underlined.

**Table S3: Individual *att*B sites for site-specific recombination systems developed to genetically engineered *Streptomyces* chromosome**

<table>
<tr><th>Name</th><th>attB sequence<br>(5′-3′)</th><th>Source of the<br>attB sequence</th><th>Ref.</th></tr>
<tr><td>µ1/6</td><td>TCTTGTAAAGCGGGGGTCGTCGGTTCGAA<br>CCCGACAGGGGGCTCAG</td><td>Streptomyces aureofaciens B96</td><td>(Farkašovská and Godány 2012)</td></tr>
<tr><td>PhiBT1</td><td>CCAGGTTTTTGACGAAAGTGATCCAGATG<br>ATCCAGC</td><td>Streptomyces coelicolor M145</td><td>(Gregory et al. 2003)</td></tr>
<tr><td>PhiC31</td><td>GGTGCCAGGGCGTGCCCTTGGGCTCCCCG<br>GGCGCG</td><td>Streptomyces lividans</td><td>(Groth et al. 2000)</td></tr>
<tr><td>PhiJoe</td><td rowspan="3">ATCTGGATGTGGGTGTCCATCTGCGGGCA<br>GACGCCGCAGTCGAAGCACGG</td><td rowspan="3">Streptomyces venezuelae ATCC 10712</td><td>(Fogg et al. 2017)</td></tr>
<tr><td>PhiOZJ</td><td rowspan="2">(Ko et al. 2020)</td></tr>
<tr><td>PhiWTR</td></tr>
<tr><td>pSAM2</td><td>GCGCGCTTCGTTCGGGACGAAGAGGT</td><td>Streptomyces ambofaciens DSM 40697</td><td>(Boccard et al. 1989; Raynal et al. 1998)</td></tr>
<tr><td>R4</td><td>AGTTGCCCATGACCATGCCGAAGCAGTGG<br>TAGAAGGGCACCGGCAGACAC</td><td>Streptomyces parvulus 2297</td><td>(Miura et al. 2011)</td></tr>
<tr><td>Samy</td><td>GAAGGTGATGCCGCGGAACTCGGGTGTG<br>TCGACGGTGCGGGTGGTGACCG</td><td>Streptomyces ambofaciens</td><td>This study</td></tr>
</table>

| | | | |
|---|---|---|---|
| | | DSM 40697 | |
| **SV1** | CATCAGGGCGGTCAGGCCGTAGATGTGGAAGAACGGCAGCACGGCGAGGACG | *Streptomyces coelicolor* A3(2) | (Fayed et al. 2014) |
| **TG1** | GATCAGCTCCGCGGGCAAGACCTTCTCCTTCACGGGGTGGAAGGTC | *Streptomyces avermitilis* ATCC 31267 | (Morita et al. 2009) |
| **VWB** | CTCTCCTAAAGCGGGTGTCGCAGGTTCGAATCCTGCCGGGGGCAC | *Streptomyces venezuelae* ETH14630 | (Van Mellaert et al. 1998) |
